# Co-opted Genes of Algal Origin Protect *C. elegans* against Cyanogenic Toxins

**DOI:** 10.1101/2022.07.06.498920

**Authors:** Bingying Wang, Taruna Pandey, Yong Long, Sofia E. Delgado-Rodriguez, Matthew D. Daugherty, Dengke K. Ma

## Abstract

Amygdalin is a cyanogenic glycoside widely used by many plants in herbivore defense. Poisonous to most animals, amygdalin-derived cyanide is detoxified by potent enzymes commonly found in bacteria and plants but not most animals. Here we show that the nematode *C. elegans* can detoxify amygdalin by a genetic pathway comprising *cysl-1, egl-9, hif-1* and *cysl-2*. Essential for amygdalin resistance, *cysl-1* encodes a protein similar to cysteine synthetic enzymes in bacteria and plants, but functionally co-opted in *C. elegans*. We identify exclusively HIF-activating *egl-9* mutations in a *cysl-1* suppressor screen and show that *cysl-1* confers amygdalin resistance by regulating HIF-1-dependent *cysl-2* transcription to protect against amygdalin toxicity. Phylogenetic analysis suggests *cysl-1* and *cysl-2* were likely acquired from green algae through horizontal gene transfer (HGT) and functionally co-opted in protection against amygdalin. Our studies reveal that HGT-mediated evolutionary changes can facilitate host survival and adaptation to adverse environment stresses and biogenic toxins.

## INTRODUCTION

In nature, many species of plants have evolved the ability to biosynthesize a group of chemically related molecules called cyanogenic glycosides as chemical toxins used in herbivore defense^1–3^. Amygdalin is a natural cyanogenic glycoside enriched in tissues of many edible plants, including seeds of stone fruits such as cherry (*Prunus avium*), peach (*Prunus persica*), apricot (*Prunus armeniaca*), and apples (*Malus domestica*). These plant species synthesize amygdalin in defense against herbivore animals as amygdalin generates poisonous cyanide upon plant tissue destruction^1–4^. After consumption by herbivores, amygdalin is hydrolyzed by β-glucosidases and α-hydroxynitrilases, releasing hydrogen cyanide or cyanogenic intermediates that can inhibit mitochondrial respiration by binding to cytochrome C oxidases, thus be highly toxic and even lethal to herbivore animals^5,6^. To tolerate amygdalin, plants also evolved a family of cysteine synthase-like enzyme, β-cyanoalanine synthases, to directly detoxify cyanides^4,7,8^. Such β-cyanoalanine synthase-like proteins are widespread and evolutionarily conserved in bacteria and plants, yet absent in most animals^8^.

*Caenorhabditis elegans (C. elegans)* is a nematode that lives with rotten fruits and bacteria in their natural ecological niches^9–11^, thus may frequently encounter amygdalin. Although this commonly used genetic model organism rarely encounters amygdalin under laboratory conditions, loss-of-function mutations in the gene *egl-9* have been shown to confer resistance to high doses of hydrogen cyanides generated by a pathogenic bacterial strain of *Pseudomonas aeruginosa*^12^. EGL-9 is a prolyl hydroxylase and negative regulator of HIF-1 (hypoxia inducible factor), a transcription factor that directly activates expression of many genes involved in hypoxic adaptation, metabolic reprogramming and stress responses^13–15^. As a transcriptional target of HIF-1, CYSL-2 is a protein strongly up-regulated in *egl-9* loss-of-function mutants and can detoxify cyanides *in vitro* as a cyanoalanine synthase^16^. CYSL-1 is a paralog of CYSL-2, both of which share sequence similarities with a family of β-cyanoalanine synthases found in bacteria and plants. We previously found that CYSL-1 regulates *cysl-2* transcription by binding to EGL-9 and promotes HIF-1 activity^17^. Whether this pathway is involved in response to cyanogenic amygdalin has not been characterized. While EGL-9 and HIF-1 are evolutionarily conserved in metazoans, most animals do not have proteins homologous to CYSL-1 and CYSL-2, except certain herbivorous arthropods^18^. In addition, co-opted detoxification genes of plant origin in whiteflies can neutralize plant-derived phenolic glycoside toxins^19,20^. However, the evolutionary origin, history of domestication and co-option of *cysl* genes in *C. elegans* remain hitherto unknown.

By screening a natural product library, we identified amygdalin as a potent activator of *C. elegans cysl-2*. We showed that the CYSL-1/EGL-9/HIF-1/CYSL-2 pathway plays a crucial role in conferring resistance to amygdalin. The intriguing acquisition of amygdalin resistance in *C. elegans* through two genes (*cysl-1* and *cysl-2*) not commonly found in other animals into the conserved EGL-9/HIF-1 genetic pathway led us to further explore the evolutionary origin and path of *cysl-1* and *cysl-2* based on phylogenetic analysis. Our combined functional and phylogenetic studies indicate that CYSL-1 and CYSL-2 may have originated from green algae by a horizontal gene transfer event in ancient nematodes and evolved by gene co-option into separate roles in a cellular circuit to respond to and detoxify cyanogenic cyanides, including amygdalin from plants today.

## RESULTS

We previously discovered several components of the HIF-1 pathway acting upstream of the oxygen-sensing HIF hydroxylase EGL-9 in *C. elegans*^17^. To identify small-molecule natural products that may modulate HIF-1 activity independently of oxygen-sensing in *C. elegans*, we conducted a natural product library screen for compounds that can activate a HIF-1-dependent transgenic reporter strain *cysl-2*p::GFP in liquid cultures under normoxia (Figure 1A). Among the identified compounds, a glycoside called amygdalin emerged as a potent activator of *cysl-2*p::GFP. We validated the finding using amygdalin from new preparations of different sources (vendor VWR or Sigma-Aldrich) to treat *C. elegans* on solid NGM media and confirmed that amygdalin can robustly activate *cysl-2*p::GFP under normoxic conditions, without affecting the transgenic co-injection marker *myo-2*p::mCherry (Figure 1B). Activation of *cysl-2*p::GFP by amygdalin is also dose-dependent and reaches peak levels at approximately 48 hrs post amygdalin treatment (Figures 1C and 1D).

**Figure 1.**
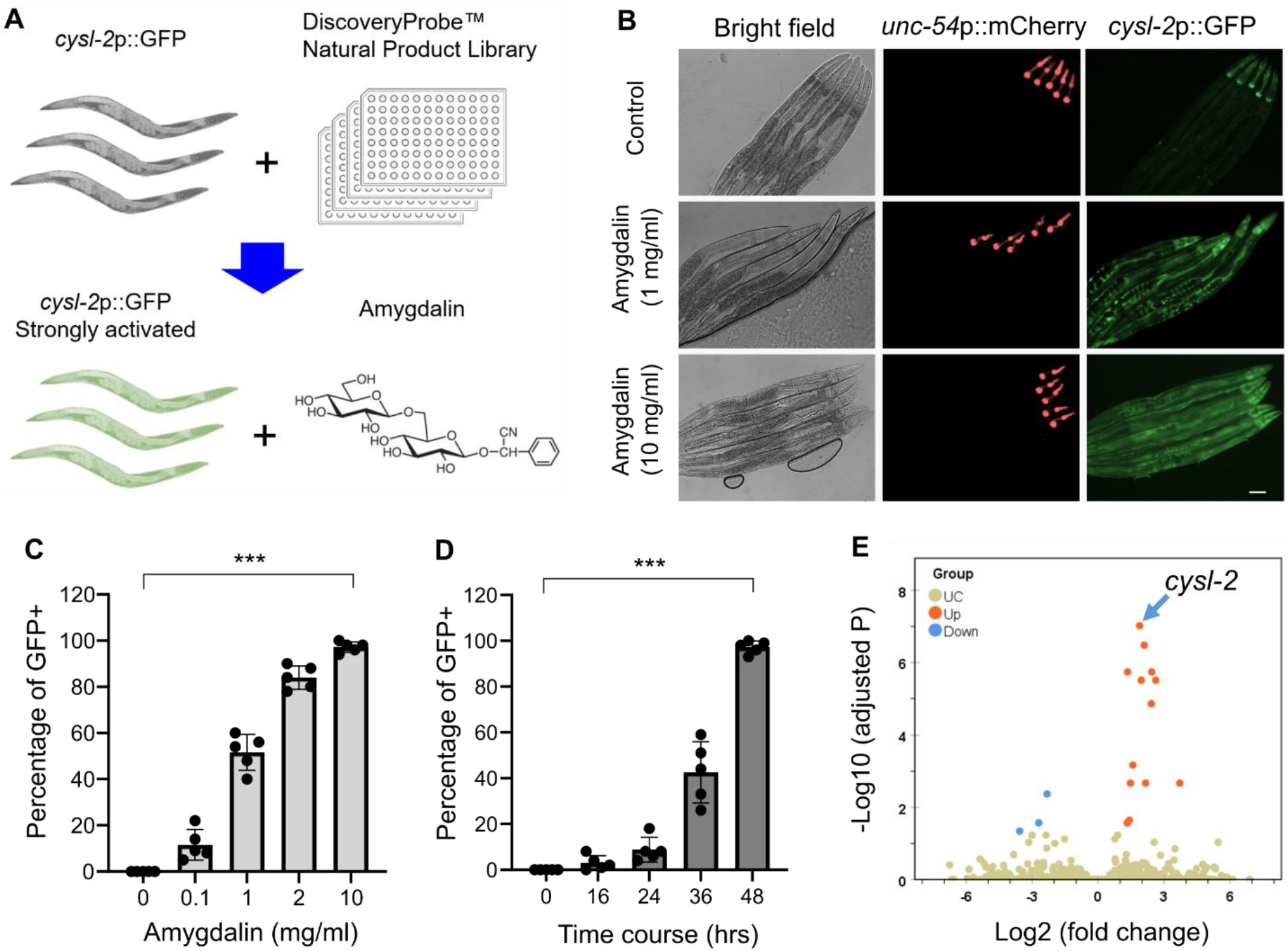
A compound screen of natural product library identifies amygdalin as a potent activator of *cysl-2*p::GFP. **(A)**, Schematic of compound screens that led to identification of amygdalin. **(B)**, Representative bright field and epifluorescence images showing dose-dependent up-regulation of *cysl-2*p::GFP by amygdalin. Scale bar: 50 µm. **(C)**, Quantification of percentages of animals with *cysl-2*p::GFP expression after 48 hrs of amygdalin treatment with increasing doses. Values are means ± S.D with ***P < 0.001 (N = 5 independent experiments, n > 50 animals per experiment). **(D)**, Quantification of percentages of animals with *cysl-2*p::GFP expression after increasing durations of amygdalin treatment at the fixed dose of 2 mg/ml. Values are means ± S.D with ***P < 0.001 (N = 5 independent experiments, n > 50 animals per experiment). **(E)**, Volcano plot showing significantly (adjusted p value < 0.05) up-(green) or down-(blue) regulated genes in amygdalin-treated animals, with *cysl-2* noted (arrow).

We performed RNAseq to identify genome-wide transcriptomic changes induced by amygdalin. We treated wild-type mixed-stage *C. elegans* hermaphrodites cultivated at 20 °C with 2 mg/ml amygdalin for 48 hrs. After differential expression analyses of triplicate samples, we identified many genes that are significantly (adjusted p value < 0.05) up-or down-regulated by amygdalin treatment (Figure 1E). As expected, *cysl-2* is among the most highly up-regulated genes (log2fold change = 2.62, adjusted p value = 3.11E-06), while expression of its upstream regulators including *hif-1, vhl-1* or *egl-9* remains largely unchanged (Figure S1 and Table S1). Gene Ontology analysis (Wormbase) indicates that amygdalin also up-regulated several genes encoding glutathione S-transferases involved in the xenobiotic detoxification pathway (Figure S1 and Table S1).

We next examined how amygdalin activates *cysl-2*p::GFP. HIF-1 and CYSL-1 are two previously established positive regulators of *cysl-2p*::GFP^17,21–23^. We found that loss-of-function mutations of *hif-1* or *cysl-1* abolished *cysl-2*p::GFP induction by low-dose amygdalin (Figure 2A). Amygdalin is a glycoside that can be metabolized sequentially to prunasin, glucose, benzaldehyde and hydrogen cyanide (Figure 2B)^1,4,5^. We treated *cysl-2*p::GFP animals with each of these amygdalin-derived molecules and found prunasin and cyanide (as in potassium cyanide) activated *cysl-2*p::GFP as amygdalin did, but others had no detectable effect (Figure 2C). CYSL-1 and CYSL-2 are cysteine synthase-like proteins and have been implicated in hydrogen sulfide and cyanide detoxification^16,17,22,24,25^. While CYSL-2 can enzymatically convert cyanide to nontoxic β-cyanoalanine^16^, the essential role of CYSL-1 in *cysl-2*p::GFP activation by amygdalin suggests CYSL-1 acts as a transcriptional regulator rather than a cysteine synthase or an enzyme to detoxify cyanide^17^. Taking advantage of the *cysl-2*p::GFP phenotype and lethal effects of amygdalin in *cysl-1* mutants (see below), we performed a *cysl-1* suppressor screen and identified exclusively three loss-of-function mutations in *egl-9* that led to constitutive *cysl-2*p::GFP expression without amygdalin treatment (Figure 2D). In further genetic epistasis analysis, loss of *hif-1* suppressed not only *cysl-2*p::GFP induction by amygdalin but also constitutive *cysl-2*p::GFP expression in *egl-9*; *cysl-1* mutants (Figure 2E). These data indicate that amygdalin activates *cysl-2*p::GFP through a dis-inhibitory regulatory pathway from cyanide, CYSL-1, EGL-9 to HIF-1 (Figure 2F).

**Figure 2.**
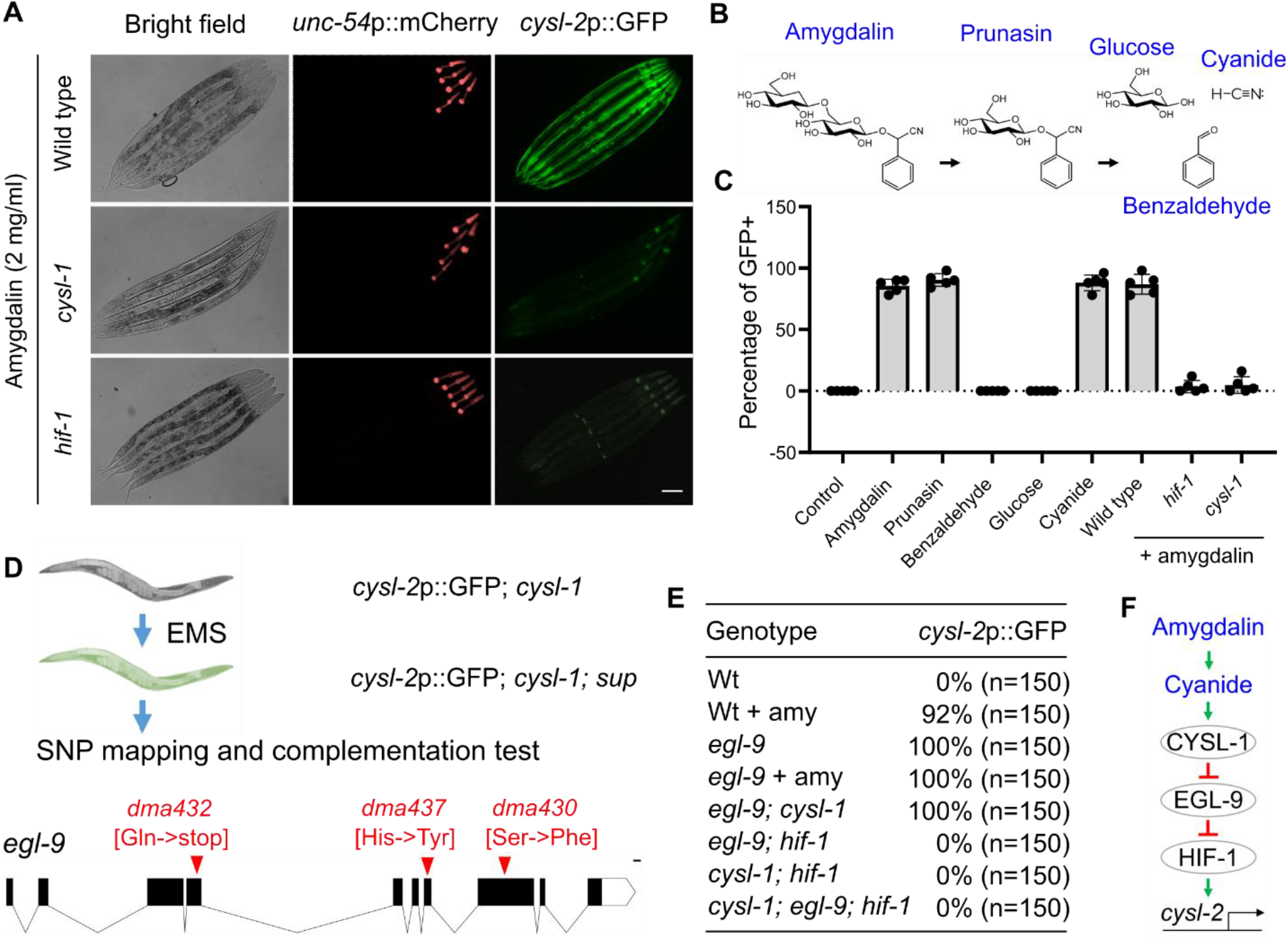
Amygdalin activates *cysl-2*p::GFP through cyanide, CYSL-1, EGL-9 and HIF-1. **(A)**, Representative bright field and epifluorescence images showing up-regulation of *cysl-2*p::GFP by amygdalin (1 mg/ml) in wild-type but not *hif-1*(*ia4*), *cysl-1*(*ok762*) loss-of-function mutant animals. Scale bar: 50 µm. **(B)**, Schematic illustrating the metabolic pathway of amygdalin, leading to generation of prunasin, glucose, hydrogen cyanide and benzaldehyde. **(C)**, Quantification of percentages of animals with strong *cysl-2*p::GFP expression after 48 hrs of treatment with amygdalin (2 mg/ml), prunacin (2 mg/ml), glucose (2 mg/ml), potassium cyanide (0.2 mg/ml) and benzaldehyde (1 mg/ml). Values are means ± S.D with ***P < 0.001 (N = 5 independent experiments, n > 50 animals per experiment). **(D)**, Schematic showing *cysl-1* suppressor screens resulting in 3 mutants, all of which are allelic to *egl-9* based on whole-genome sequencing and complementation tests. **(E)**, Table summary of *cysl-2*p::GFP activation in animals with indicated genotypes and treatment conditions. Penetrance and numbers of animals examined are noted. **(F)**, Model illustrating the regulatory pathway that mediates the transcriptional response to amygdalin.

To assess the functional importance of the CYSL-1/EGL-9/HIF-1/CYSL-2 pathway, we examined the time-dependent survival response to high-dose amygdalin (10 mg/ml on NGM; effective concentrations *in vivo* are likely much lower owing to cuticle and intestinal absorption barriers) in animals with individual or combinatorial pathway gene mutations. Wild-type animals exhibited no apparent decrease in survival upon amygdalin over three days of treatment (Figure 3A). By contrast, loss-of-function *cysl-1* mutants displayed striking sensitivity to amygdalin, with nearly complete population death at day 3 of amygdalin treatment (Figure 3B). Loss-of-function of *cysl-2* also caused similar time-dependent sensitivity to amygdalin (Figure 3C). While loss of *egl-9* fully suppressed vulnerability of *cysl-1* mutants to amygdalin (Figure 3D), loss of either *hif-1* or *cysl-2* rendered *egl-9* mutants sensitive to amygdalin (Figures 3E and 3F). These data support central roles of the CYSL-1/EGL-9/HIF-1/CYSL-2 pathway in the transcriptional response and conferring resistance to amygdalin in *C. elegans*.

**Figure 3.**
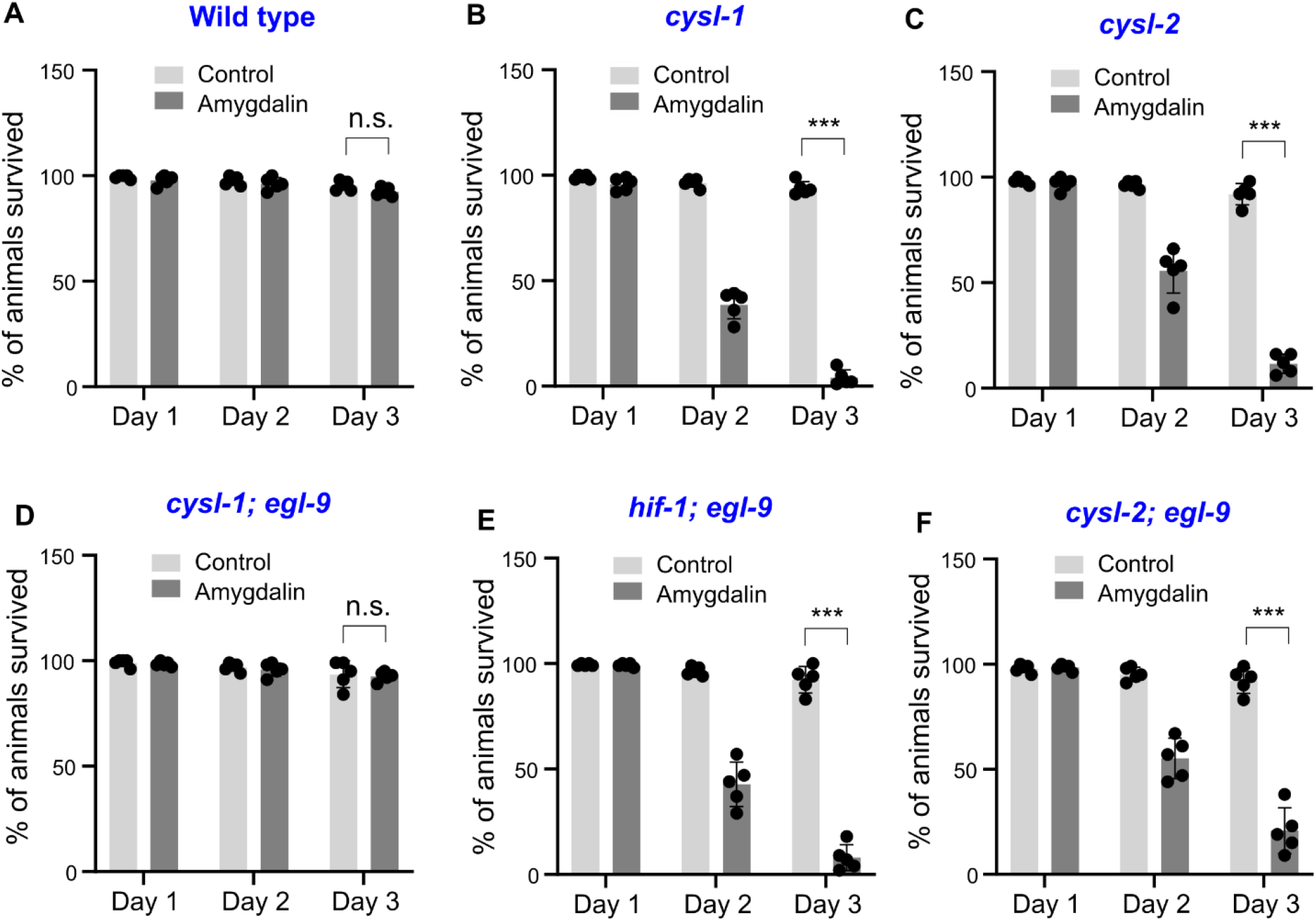
Amygdalin resistance requires CYSL-1, HIF-1 and CYSL-2 acting in a regulatory pathway. **(A)**, Quantification of percentages of post-L4 stage wild-type animals survived after 24, 48 or 72 hrs of treatment with amygdalin (10 mg/ml). **(B)**, Quantification of percentages of post-L4 stage *cysl-1*(*ok762)* loss-of-function mutant animals survived after 24, 48 or 72 hrs of treatment with amygdalin (10 mg/ml). **(C)**, Quantification of percentages of post-L4 stage *cysl-2*(*ok3516)* loss-of-function mutant animals survived after 24, 48 or 72 hrs of treatment with amygdalin (10 mg/ml). **(D)**, Quantification of percentages of post-L4 stage *cysl-1*(*ok762); egl-9(sa307)* double loss-of-function mutant animals survived after 24, 48 or 72 hrs of treatment with amygdalin (10 mg/ml). **(E)**, Quantification of percentages of post-L4 stage *hif-1*(*ia04); egl-9(sa307)* double loss-of-function mutant animals survived after 24, 48 or 72 hrs of treatment with amygdalin (10 mg/ml). **(F)**, Quantification of percentages of post-L4 stage *cysl-2*(*ok3516); egl-9(sa307)* double loss-of-function mutant animals survived after 24, 48 or 72 hrs of treatment with amygdalin (10 mg/ml). Values are means ± S.D with ***P < 0.001 (N = 5 independent experiments, n > 50 animals per experiment).

To investigate the evolutionary origin *cysl-1/2* in *C. elegans*, we searched for homologs of CYSL-1 and CYSL-2 proteins across all domains of life. We identified related proteins with >50% sequence identity in nematodes, green algae, land plants, and bacteria. Outside of nematode proteins, the most closely related sequences, and the only ones with sequence identities >60% were from the *Chlorophyta* clade of green algae (Figures S2A and 4A).

**Figure 4.**
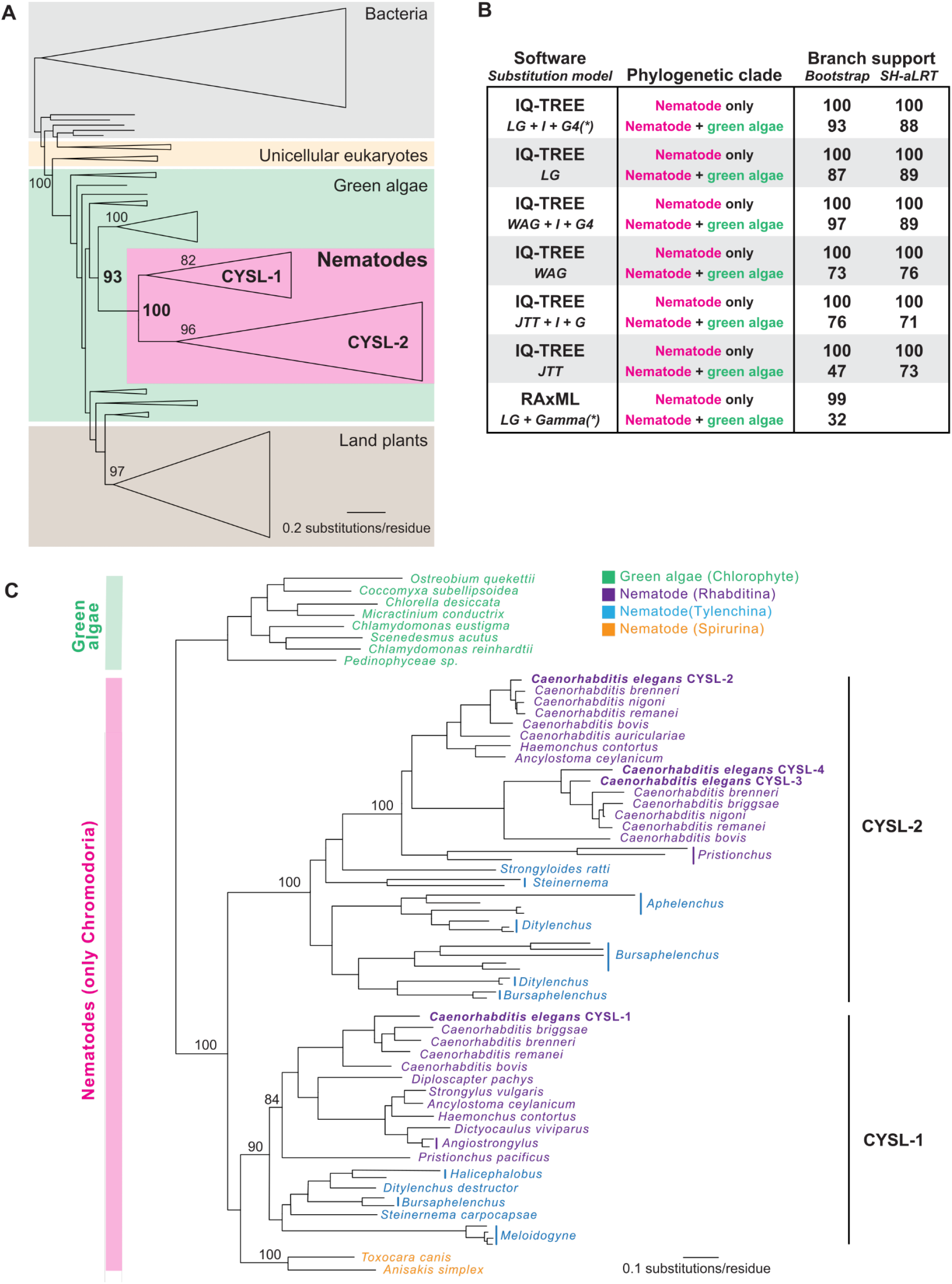
Nematode *cysl* genes were likely acquired from green algae by horizontal gene transfer. **(A)**, IQ-TREE generated maximum likelihood (ML) phylogenetic tree of homologs of nematode CYSL proteins across a broad range of species. Species groups are shaded. Clades of similar proteins are collapsed and shown as triangles. Bootstrap support values for relevant branches are shown. Bolded values indicate the support for a clade that only contains CYSL-like proteins from nematodes (100% bootstrap support) and a well-supported clade that contains only CYSL-like proteins from nematodes and a group of green algae in the *Chlorophyta* lineage (93% bootstrap support). Within the nematodes, two clear, well-supported clades of CYSL proteins exist, one that contains *C. elegans* CYSL-1 and one that contains *C. elegans* CYSL-2. **(B)**, ML phylogenetic analyses were performed with multiple programs and multiple amino acid substitution models as indicated. Support values (bootstrap and SH-aLRT when available) are shown for a clade containing only nematode proteins and a clade containing nematode and chlorophyte green algae proteins as are bolded in part A. Asterisks next to model names indicate that best fitting model as described in Methods. **(C)**, Expanded phylogenetic tree of CYSL proteins from nematodes and their nearest homologs in green algae. Font colors denote major groups of nematode species as found in^26^. Bootstrap support values are shown at relevant nodes. For panels A-C, a complete list of all sequences used is in Supplementary Data file 1 and all trees, including support values, are found in Supplementary Data file 2.

To determine the phylogenetic origin of CYSL proteins in nematodes, we performed maximum likelihood (ML) based phylogenetic analyses on CYSL homologs from a broad range of eukaryotic and prokaryotic species (Data Files S1, S2 and Figure 4A). Consistent with the high sequence similarity to proteins in green algae, we found strong support for nematode proteins nesting within the chlorophyte clade of green algae (Figure 4B). Proteins from streptophytes (*Streptophyta*), which include some green algae and all land plants, are found in a distinct region of the phylogenetic tree, as are proteins from single-cell eukaryotes (*e.g*. groups including *SAR, Excavata*, and *Rhodophyta*) and bacteria. These inferences were robust to the ML software we used for analysis, as well as the substitution models we employed (Figure 4B). Importantly, cysteine synthase enzymes found in some arthropods, such as mites, which are the result of a horizontal gene transfer (HGT) event from bacteria show less than 40% sequence identity, consistent with their separate evolutionary origin (Figure S2B)^18^. Taken together, these data support a single HGT event from a green algal species in the *Chlorophyta* that gave rise to nematode CYSL.

We further wished to analyze the distribution of CYSL proteins in nematodes to determine at what point CYSL was acquired by nematodes and at what point gene duplication occurred (Figure 4C). We found CYSL proteins only in the *Chromadoria* clade of nematodes and not in the well sampled organisms in the *Trichinellida* (*e.g. Trichinella* species). Although we cannot rule out that it was lost in the *Trichinellida*, the most parsimonious explanation for this distribution is that CYSL was acquired by an ancestral *Chromodoria* nematode after the divergence from other clades of nematodes. Within *Chromodoria*, we only detected a single CYSL homolog in the most basally branching *Spirurina* nematodes^26^ (*e.g. Toxocara canis*) whereas we observed two or more copies of CYSL homologs in *Tylenchina* (*e.g. Bursaphelenchus* species) and *Rhabditina* (*e.g. Caenorhabditis* species) lineages of nematodes. These data, coupled with the branching of CYSL-1 and CYSL-2 paralogs at the base of the nematode CYSL protein phylogeny (Figure 4C), suggest that CYSL duplicated soon after acquisition in nematodes to CYSL-1-like and CYSL-2-like proteins. Based on our findings of the non-overlapping functional importance of CYSL-1 and CYSL-2 in *C. elegans* for cyanide detoxification, these phylogenetic findings suggest that this network of CYSL-1 and CYSL-2 evolved soon after the acquisition of these enzymes by nematodes.

## DISCUSSION

In this study, we establish novel physiological roles of the CYSL-1/EGL-9/HIF-1/CYSL-2 pathway in the detoxification of amygdalin in *C. elegans*. Our data indicate that CYSL-1 acts upstream of the evolutionarily conserved HIF-1 pathway as a key regulator of *cysl-2* induction upon amygdalin exposure, resulting in detoxification of cyanide that likely derives from natural diets (e.g. cyanogenic plant tissues and *Pseudomonas* bacteria) of *C. elegans* and other nematodes. How amygdalin or cyanide is sensed by *C. elegans* and regulates CYSL-1 and EGL-9 awaits further studies. Plausible mechanisms include mitochondrial inhibition by cyanide, posttranslational modification by S-cyanylation, and allosteric modulation of CYSL-1 and EGL-9 interaction^6,17,27^.

By phylogenetic analysis, we obtained evidence pointing to an intriguing evolutionary origin of *cysl-1* and *cysl-2* in *C. elegans*. Maximum likelihood estimation based on protein sequence similarity supports a single HGT event from a green algal species in the *Chlorophyta* that gave rise to nematode *cysl* genes. Our analysis further suggests that a *cysl* gene ortholog was acquired by ancestral *Chromodoria* nematodes after the divergence from other clades of nematodes, and duplicated soon after their acquisition to give rise to *cysl* gene paralogs. In *C. elegans, cysl-1* and *cysl-2* have further evolved separate functions from ancestral cysteine synthase-like activity into division of labor. While CYSL-2 retains cyanoalanine synthase enzymatic activity to detoxify cyanide, CYSL-1 is co-opted to regulate *cysl-2* expression upon amygdalin/cyanide exposure, in an elegant cellular circuit that detects and subsequently defends against toxicity associated with amygdalin/cyanide.

*Chromodoria* nematodes diverged from other clades of nematodes more than 250 million year ago, around which drastic geological and ecological changes (e.g. those leading to Permian-Triassic mass extinction) may have exerted selection pressures on nematodes surviving in environments rife with sulfide/cyanide toxicity and ancient algal species^28–32^. Following acquisition of *cysl* genes, nematodes may have retained and co-opted *cysl* gene functions in adaptation to new environments, including living as plant parasites or in rotten fruits associated with cyanide presence. While precise mechanisms of these evolutionary changes particularly HGT remain unclear, the evolutionary pathway leading to distinct cellular roles of *cysl-1* and *cysl-2* in *C. elegans* today exemplifies how organisms can acquire novel biological traits to survive and reproduce by HGT and gene co-option^32–37^. Functional HGTs within prokaryotic organisms and from bacteria to eukaryotes that have been widely observed and characterized^38–41^. Our functional and phylogenetic analyses reveal previously unknown HGT from algae to nematodes, supporting the emerging theme that such events of HGTs can lead to evolutionary changes and novel traits to facilitate host protection and adaptation to abiotic stress, chemical toxins and extreme environments.

## Figures and Figure legends

**Figure S1.**
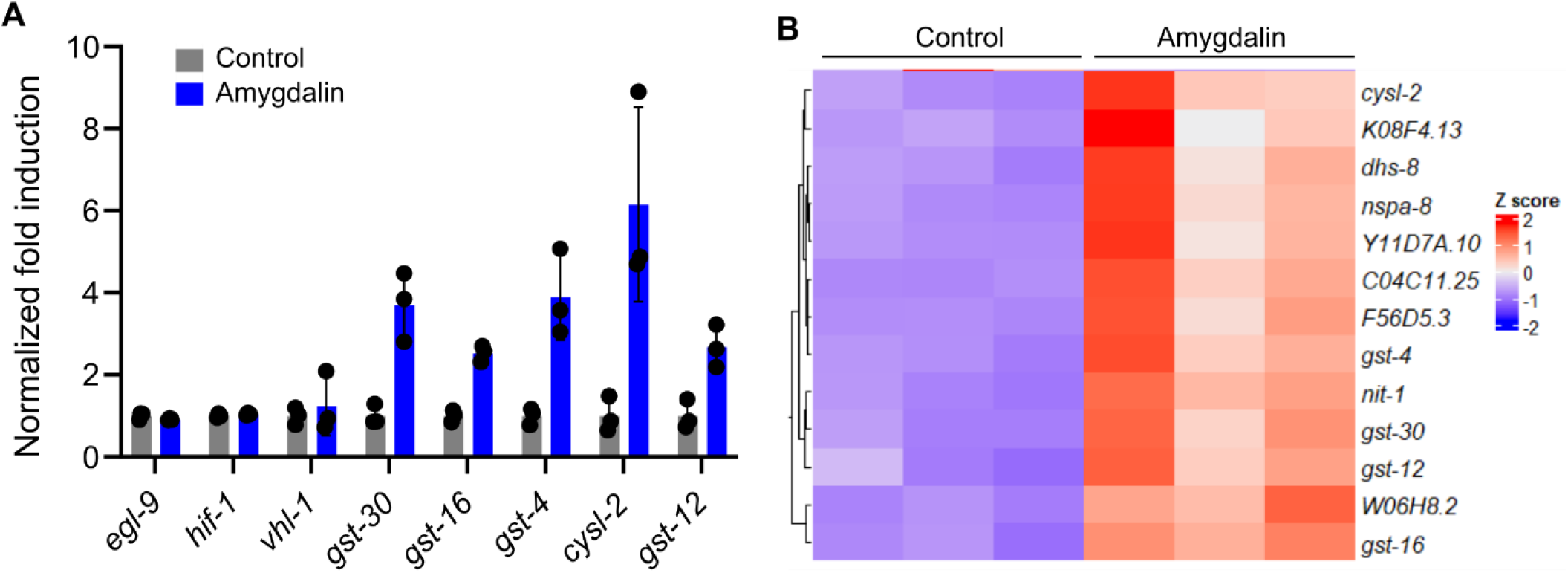
Amygdalin-independent and amygdalin-regulated genes identified by RNAseq involved in the HIF and xenobiotic detoxification pathways. **(A)**, Normalized fold induction by amygdalin (2 mg/ml on NGM for 48 hrs) of genes acting upstream of *cysl-2* (*egl-9, hif-1, vhl-1*) and GST-encoding genes involved in xenobiotic detoxification. **(B)**, Heat map showing hierarchical clustering of genes differentially up-regulated by amygdalin (2 mg/ml on NGM for 48 hrs) from RNAseq experiments.

**Figure S2.**
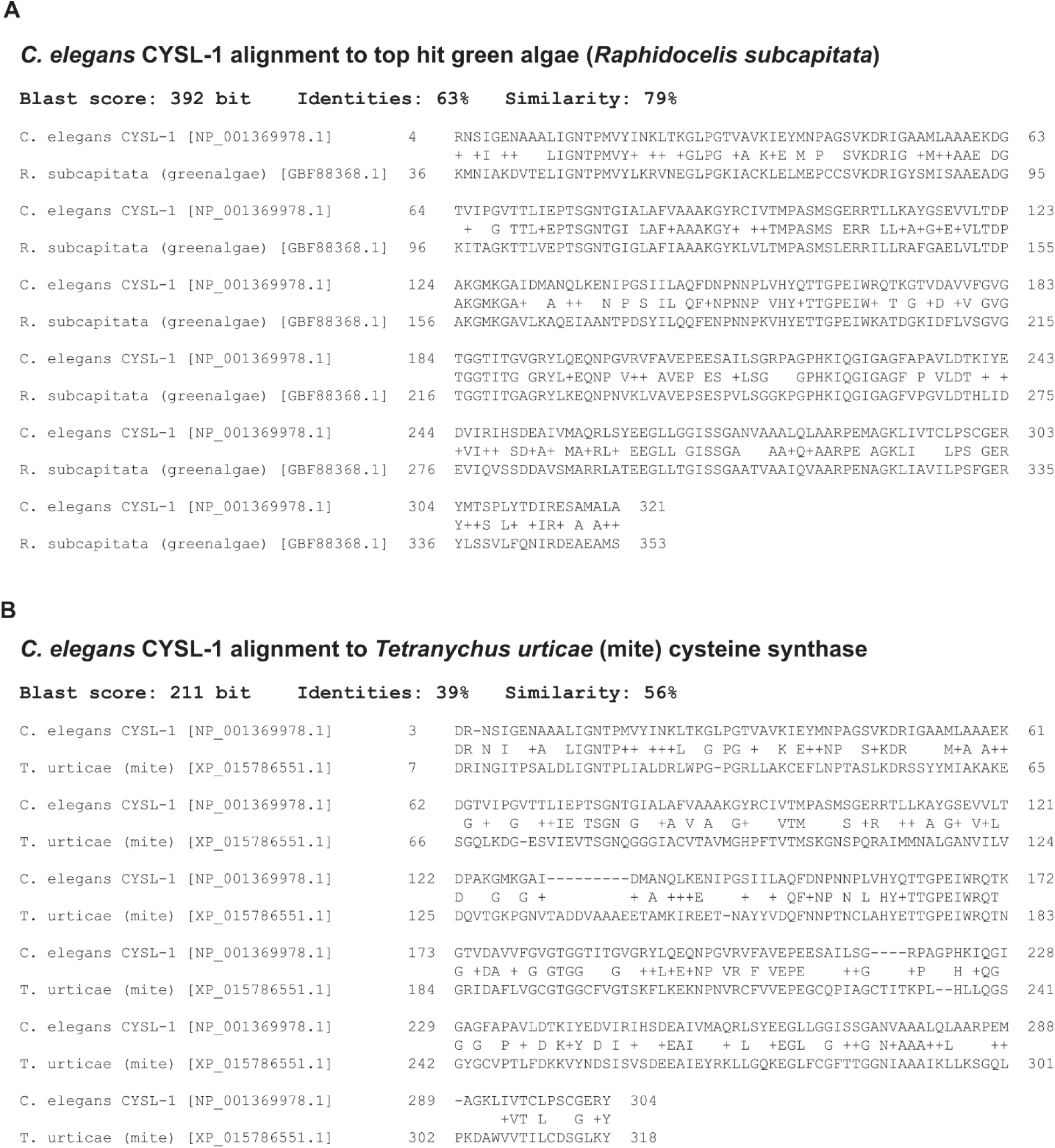
Protein sequence alignments of CYSL-1 with orthologs from green algae (A) and mite (B). Blast scores, a.a. identities and similarities are shown.

## Methods and Materials

### C. elegans

*C. elegans* strains were maintained under standard laboratory conditions unless otherwise specified. The N2 Bristol strain was used as the reference wild type, and the polymorphic Hawaiian strain CB4856 was used for genetic linkage mapping and SNP analysis ^42,43^. Forward genetic screens for *cysl-2*p::GFP; *cysl-1* suppressing mutants after ethyl methanesulfonate (EMS)-induced random mutagenesis were performed as described previously ^17,44^. Identification of mutations by whole-genome sequencing and complementation tests by crossing EMS mutants with *egl-9(sa307)* heterozygous males were used to determine *dma430, dma432* and *dma437* as new loss-of-function alleles of *egl-9*. Transgenic strains were generated by germline transformation ^45^. Transgenic constructs were co-injected (at 25 ng/μl) with *unc-54p*::mCherry, and extrachromosomal lines of fluorescent animals were established. Genotypes of strains used are as follows: Chr. II: *cysl-2*(*ok3516*); Chr. IV: *nIs470 [cysl-2p::GFP; unc-54p::mCherry]*, Chr. V: *egl-9(dma430, 432, 437, sa307), hif-1(ia4)*, Chr. X: *cysl-1(ok762)*.

### Sample preparation for RNA sequencing and data analysis

N2 control (M9 treated) and amygdalin-treated animals (2 mg/ml on NGM for 48 hrs) were maintained at 20 °C and washed down from NGM plates using M9 solution and subjected to RNA extraction using the RNeasy Mini Kit from Qiagen. RNA-seq library preparation and data analysis were performed as previously described ^46^. Three biological replicates were included for each treatment. The cleaned RNAseq reads were mapped to the genome sequence of *C. elegans* using the hisat2 tool^47^. Abundance of genes was expressed as FPKM (Reads per kilobase per million mapped reads). Identification of differentially expressed genes was performed using the DESeq2 package ^48^.

### *C. elegans* compound screen, amygdalin sensitivity and survival assays

*C. elegans* animals carrying the integrated *cysl-2*p::GFP reporter were cultured under non-starved conditions for at least 4 generations before natural product screens using the DiscoveryProbe Bioactive Compound Library (Catalog No. L1022P, APExBIO). Synchronized L1-stage animals were grown to the L4 stage and transferred to deep 96-well plates in liquid cultures containing each compound (0.5 mM in 100 µL volume, approximately 100 animals per well) in S medium with concentrated *E. coli* OP50 as a food source. The cultures were incubated at 20 °C for 48 hrs before observation under an epifluorescence microscope (SMZ18). Candidate hits were subject to multiple trials and repeated with the same compounds from different providers and to determine phenotypic penetrance, fluorescence intensity and compound dose-dependency.

*C. elegans* animals were cultured under non-starved conditions for at least 4 generations before amygdalin sensitivity assays. NGM plates spread with equal agar thickness seeded with equal amounts of freshly seeded OP50 (within 24 hrs of seeding) were used for M9 or amygdalin (dissolved in M9, TCA0443-010G, VWR or equivalent A6005, Sigma-Aldrich) supplementation. Fresh preparations of fully dissolved amygdalin in M9 were added (250 µl/ 60 cm plate) and spread evenly on NGM plates with pre-seeded OP50. Once the plates with amygdalin were briefly air dried, synchronized-stage (L4) animals were placed and monitored for survival over 24, 48 and 72 hrs post treatment. Animals were scored as dead if they showed no pumping and movement upon light touch with the body necrosis subsequently confirmed.

### Phylogenetic analysis

*C. elegans* CYSL-1 (accession NP_001369978.1) was used as a BLASTp ^49^ search query against the nonredundant (NR) and Reference Sequence (RefSeq) protein databases. Resulting sequences were aligned using Clustal Omega ^50^ using two iterations of refinement and duplicate sequences were removed. Non-aligning termini and alignment positions in which >95% of all sequences had a gap were manually removed from subsequent analyses. To generate phylogenetic trees encompassing a broad range of CYSL homologs, sequences with >80% sequence identity were reduced to a single unique sequence using CD-HIT ^51^ with a 0.8 sequence identity cutoff. The resulting 253 protein sequences (accession numbers and species names found in Supplemental file 1) were realigned as above and used as input for maximum likelihood phylogenetic programs. IQ-TREE ^52^ phylogenies were generated using the “-bb 1000 -alrt 1000” commands for generation of 1000 ultrafast bootstrap ^53^ and SH-aLRT support values. The best substitution model was determined by ModelFinder ^54^ using the “-m AUTO” command or the substitution model was specified as shown in Figure 4B using the “-m” command. RAxML phylogenies were generated using “-f a” with 500 bootstrap replicates and “-m PROTGAMMAAUTO” to determine the best substitution model with gamma distribution. All phylogenetic trees generated from IQ-TREE and RAxML, with support values, can be found in Supplemental file 2.

To generate a phylogeny of nematode CYSLs, the relevant subtree containing nematodes and the nearest green algae was extracted for the larger phylogeny generated above. Additional nematode sequences that had been removed using the CD-HIT filtering described above were added back in. Sequences were realigned and degapped as described above with a CDHIT similarity cutoff of 98% followed by additional manual removal of poorly aligning sequences. An IQ-TREE phylogenetic tree using ModelFinder (LG+I+G) was determined as described above. Accession numbers and species names are found in Supplementary Data file 1, and complete IQ-TREE with support values is found in Supplementary Data file 2.

### Confocal and epifluorescence microscopic imaging

SPE epifluorescence compound microscopes (Leica) were used to capture fluorescence images. Animals of different genotypes were randomly picked at the same young adult stage (24 hrs post L4) and treated with 1 mM Levamisole sodium Azide in M9 solution (31,742-250MG, Sigma-Aldrich), aligned on an 4% agar pad on slides for imaging. Identical setting and conditions were used to compare genotypes, treatment experimental groups with control.

### Statistical analysis (wet lab data)

Data were analyzed using GraphPad Prism 9.2.0 Software (Graphpad, San Diego, CA) and presented as means ± S.D. unless otherwise specified, with *P* values calculated by unpaired two-tailed t-tests (comparisons between two groups), one-way ANOVA (comparisons across more than two groups) and adjusted with Bonferroni’s corrections.

## Data availability

The RNAseq read datasets were deposited in NCBI SRA (Sequence Read Archive) under the BioProject accession PRJNA843348. All other data generated for this study are included in this article.

## Acknowledgments

Some strains were provided by the CGC, which is funded by NIH Office of Research Infrastructure Programs (P40 OD010440). The work was supported by NIH grant 1R35GM139618 (D.K.M.), Packard Fellowships for Science and Engineering (D.K.M.) and the Pew Scholars Program in the Biomedical Sciences (D.K.M. and M.D.D.).

## Author contributions

B.W., T.P. and D.K.M. conceptualized, performed and analyzed the wet experiments and wrote the manuscript. Y.L. performed RNAseq bioinformatic analysis. S.D.R. and M.D.D. performed phylogenetic analysis and wrote the manuscript. M.D.D and D.K.M. supervised the project.

